## Supplemental Figures for "Upregulated Ca^2+^ release from the endoplasmic reticulum leads to impaired presynaptic function in Alzheimer’s disease"

### Supplementary Figures

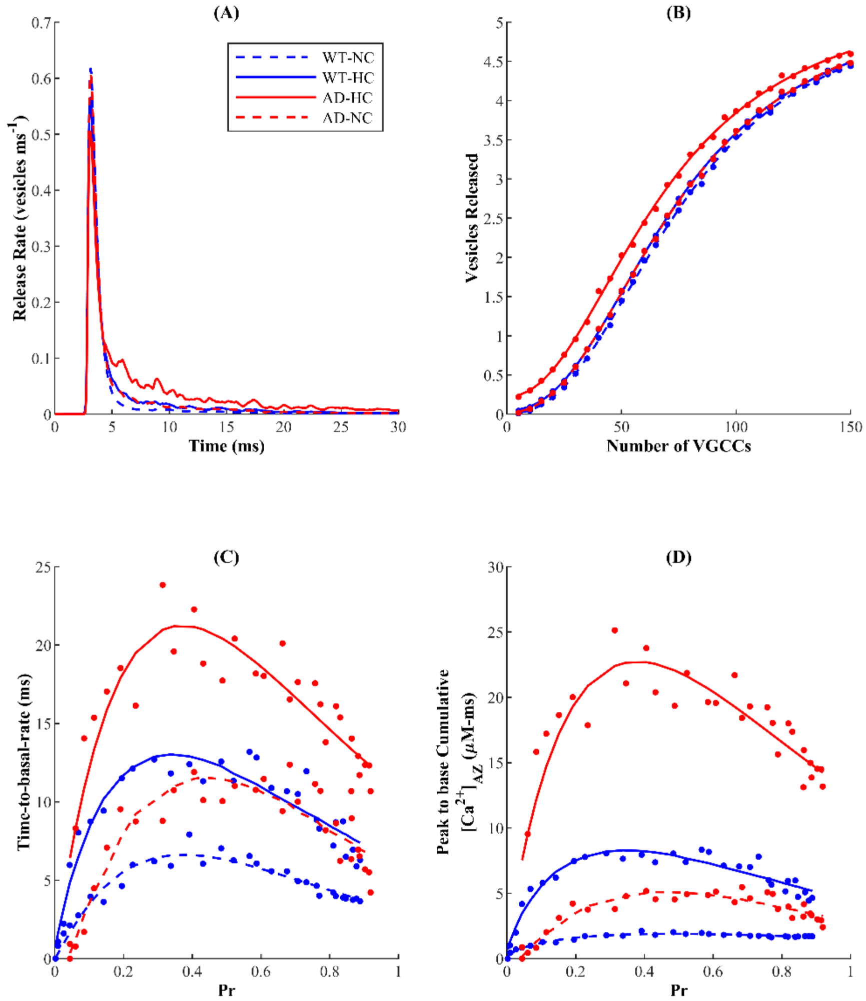


**SI Figure 1.** *Coupling between the microdomains around $IP_3Rs$ and VGCCs influences $Ca^{2+}$ in the AZ and neurotransmission profiles in AD and WT synapses*. Transmitter release rates within 30 ms of stimulus (A) and total vesicles released (B) for different coupling configurations. Decay time of peak release rate (C) and cumulative $Ca^{2+}$ concentration from peak to basal rate in the AZ (D) are markedly influenced by coupling.

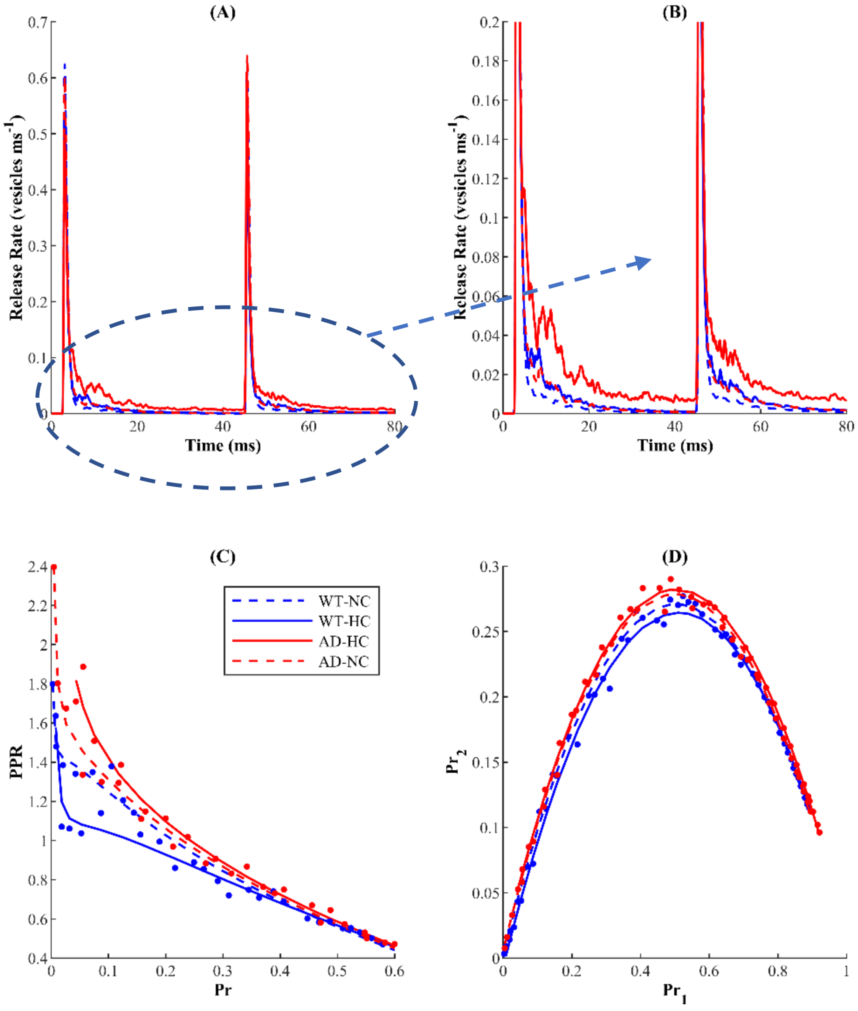


**SI Figure 2.** *Stronger coupling between the microdomian of $IP_3Rs$ cluster and AZ exacerbate the release rate and enhanced PPR in AD-affected synapses but only marginally affect the bell-shaped behavior of $Pr_2$ as a function of $Pr_1$***.** Release profile (A) (zoomed-in (B)) following paired-pulse stimulation protocol shows an increase in release rate by stronger coupling in the microdomian of $IP_3Rs$ cluster and AZ in both WT and AD-affected synapses. (C) The enhanced PPR in AD-affected synapses with respect to WT synapses is exacerbated by stronger coupling. (D) Pr in response to the second pulse ($Pr_2$) as a function of Pr following the first pulse ($Pr_1$) shows that the bell-shaped response is marginally affected by the coupling strength.

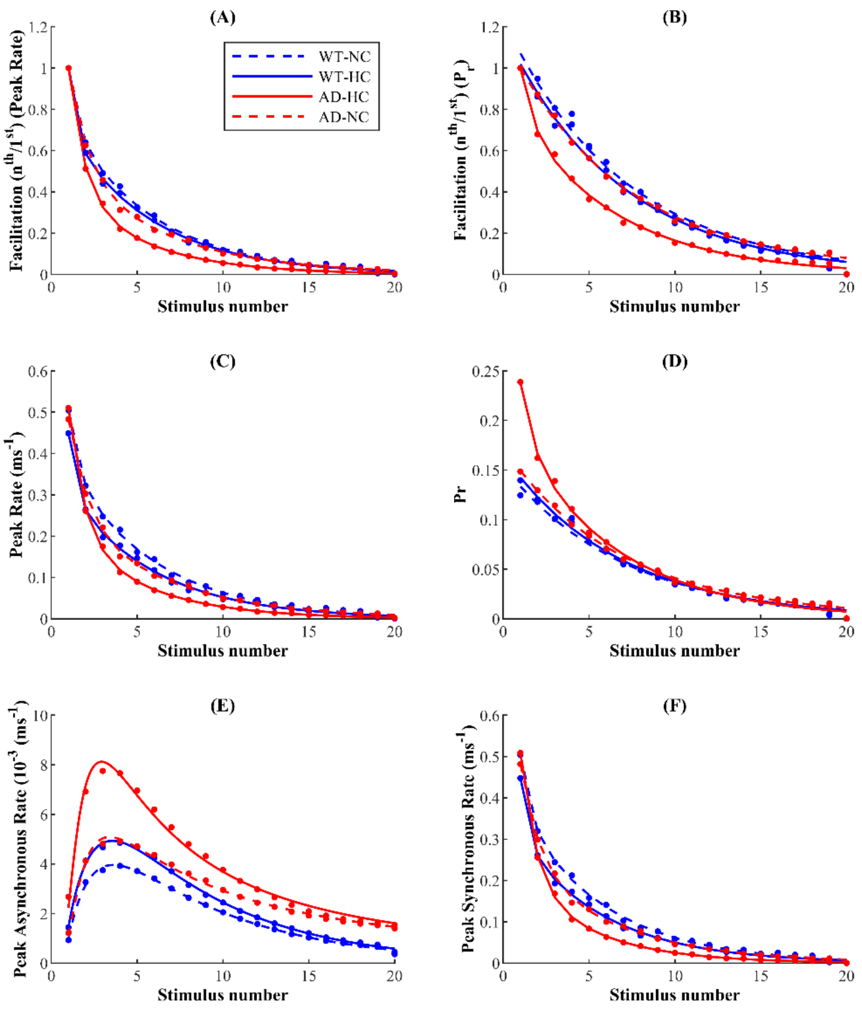


**SI Figure 3.** *Stronger coupling between the microdomian of $IP_3Rs$ cluster and AZ exacerbate the stronger depression in AD-affected synapses*. Facilitation obtained from peak rate (A) and Pr (B) shows that HC enhances the synaptic depression in AD-affected synapses. Peak release rate (C) and Pr (D) following each AP in the train under different coupling conditions. (E) Asynchronous and (H) peak synchronous release under different coupling conditions.
